# Benchmarking of Optimization Algorithms for Boolean Model Inference in Biomedicine

**DOI:** 10.1101/2025.09.12.675817

**Authors:** Bingyu Jiang, Pierre Klemmer, Marek Ostaszewski

## Abstract

Biological processes in health and disease are regulated in great complexity, imposing significant challenges in understanding and modifying their behavior for healthcare applications. Boolean networks have become essential tools for modeling gene regulatory systems and understanding cellular decision-making processes, but their optimization for biological relevance and precision medicine remains challenging.

This study presents a comprehensive benchmark comparison of four prominent Boolean network optimization methods involving genetic algorithms, integer linear programming, and answer set programming, evaluating their performance across structural robustness, method reliability, and biological relevance using mean squared error (MSE) as the primary optimization criterion. Through systematic analysis of network reconstruction under varying perturbation levels (10-90%), we demonstrate that each method exhibits distinct performance profiles: answer set programming (ASP) achieves optimal topological similarity with computational efficiency, integer linear programming (ILP) produces reasonable MSE minimization but with high variance, genetic algorithms (GA) shows superior functional reconstruction stability despite longer computational times. Our results reveal critical limitations in current evaluation approaches, particularly the insufficient discriminatory power of F1 scores and Hamming distance metrics, and highlight fundamental trade-offs between data fitting accuracy and topological preservation.

The analysis demonstrates that no single optimization method dominates across all criteria, with all methods showing significant performance degradation at perturbation thresholds above 10-30%, suggesting that method selection should be application-specific and guided by requirements for computational efficiency, reconstruction accuracy, and robustness to uncertainty in prior knowledge.

## 1 Introduction

Classically, disease-relevant mechanisms are investigated in costly and time-consuming wet lab experiments, lending precise molecular insights into highly contextual interactions. Comprehensively understanding a plethora of novel biological insights is a daunting task for any researcher given the increasing mass of biomedical publications [16]. Systematic representations like network diagrams compile biomedical knowledge into curated semantic formats that ideally follow community guidelines [14], while simultaneously providing solid frameworks for computational analysis. Network and pathway diagrams can be interpreted as mathematical models representing correlational or causal relationships between bio-molecules [18], providing the ability to predict the effects of perturbations and interventions into the underlying biological systems. Systems biology approaches like modeling take holistic stances in applying computational methods to high-throughput biological data to identify patterns and regulatory chains, accurately simulating complex system behaviors on the molecular scale.

Boolean models have emerged as a powerful framework for understanding complex biological systems, particularly in the context of gene regulatory networks. Originally introduced by Kauffman [8] in the 1960s, Boolean networks have been extensively applied to model genetic regulatory systems, providing insights into cellular decision-making processes and disease mechanisms [7]. A Boolean network is a mathematical model used to represent complex systems where each component can exist in one of two states, typically denoted as 0 (inactive) or 1 (active). It is composed of nodes and edges. Nodes represent the components of the system, such as genes, proteins, or molecular species. Each nodes state evolves over time according to a logical rule. Edges represent interactions and rules between nodes, indicating how one component influences another. They typically encode activation (positive influence) or inhibition (negative influence). Node states are updated based on the states of their input nodes and the logical rules associated with the edges, allowing the network to capture dynamic behaviors such as stable states, oscillations, and signal propagation. Stable states, including cyclic oscillatory states, are referred to as attractors and are of special interest given that the system cannot escape them without external inputs. As such, they are interpreted as analogs to biological phenotypes - if a model representing the cell cycle reaches a steady state, it may be interpreted as cell cycle arrest, while a cyclic attractor represents normal cell cycle continuation.

As Boolean networks have become increasingly relevant in systems biology, numerous algorithms have been developed to optimize model structure for better biological relevance and predictive accuracy. Systems biology diagrams often describe mechanisms in overgeneralized contexts that fail to capture specific genetic or environmental variation, leading to poor applicability of such general models. Ever-increasing amounts of biomedical data make it increasingly feasible to construct personalized and structurally optimized models that provide better fits to specific patients.

The goal of this project is to investigate whether structure-based optimization using experimental data leads to functionally equivalent networks in terms of dynamical behavior. We compare four prominent optimization methods implemented in three toolkits: ASP in caspo [6], variable neighborhood search (VNS) in MEIGO [3], as well as ILP and GA in CellNOpt [5] using mean squared error (MSE) as our primary metric.

These optimization methods address a fundamental biological question: Given a proposed network structure, which parts of it are actually relevant under specific experimental conditions? Real biological networks are inherently context-dependent; the same cell might harbor hundreds of potential regulatory connections, but only a subset remain active under specific conditions such as drug treatment, disease states, or developmental stages. This context-dependency necessitates sophisticated optimization approaches that can identify the most relevant network components for given experimental scenarios.

Through systematic comparison of these methods, our investigation focuses on three critical aspects of optimization method performance:

1. **Robustness Testing:** How sensitive are these optimization methods to errors in the input network topology? This question is particularly relevant given that prior knowledge networks (PKNs) used as starting points unavoidably contain incomplete or inaccurate information about regulatory relationships.
2. **Method Reliability:** Do different algorithms converge to similar solutions when starting from slightly different network topologies? Consistency across multiple runs and minor variations in input conditions is essential for generating reliable biological insights.
3. **Biological Relevance:** Do the optimized networks capture known biological interactions and pathways? The ultimate validation of any optimization method lies in its ability to preserve biologically meaningful relationships while eliminating spurious connections.

Generally, these algorithms produce modified candidate versions of the input PKN whose topology and dynamics are compared to the supplied experimental data. If the proposed candidate reproduces the experimental observations faithfully under the same initial conditions, it is thought to be a suitable approximation of the underlying biological system. The detailed algorithmic descriptions can be found in the supplementary document (table 2 shows a summary of these methods). This essay focuses primarily on developing suitable benchmarks to compare different approaches from numerous perspectives.

**Table 1:** Dataset summaries. Experimental data structure is elaborated on further in section 2.1.1.

|  | **Toy dataset** | **DREAM dataset** | **T-Cell Data** |
| --- | --- | --- | --- |
| **Network summary** |  |  |  |
| Nodes (species) | 15 | 40 | 40 |
| Edges (reactions) | 16 | 58 | 52 |
| Activating edges | 15 | 56 | 47 |
| Inhibitory edges | 1 | 2 | 5 |
| **Experimental data summary** |  |  |  |
| Cues (inputs measured) | 4 | 8 | 6 |
| Stimuli applied | 2 | 4 | 3 |
| Inhibitors applied | 2 | 4 | 3 |
| Signals read out | 7 | 7 | 8 |
| Time-points per experiment | 2 | 2 | 2 |
| Total experiments (rows) | 9 | 25 | 17 |
| Number of Attractors | 4 | 16 | 10 |

**Table 2:**
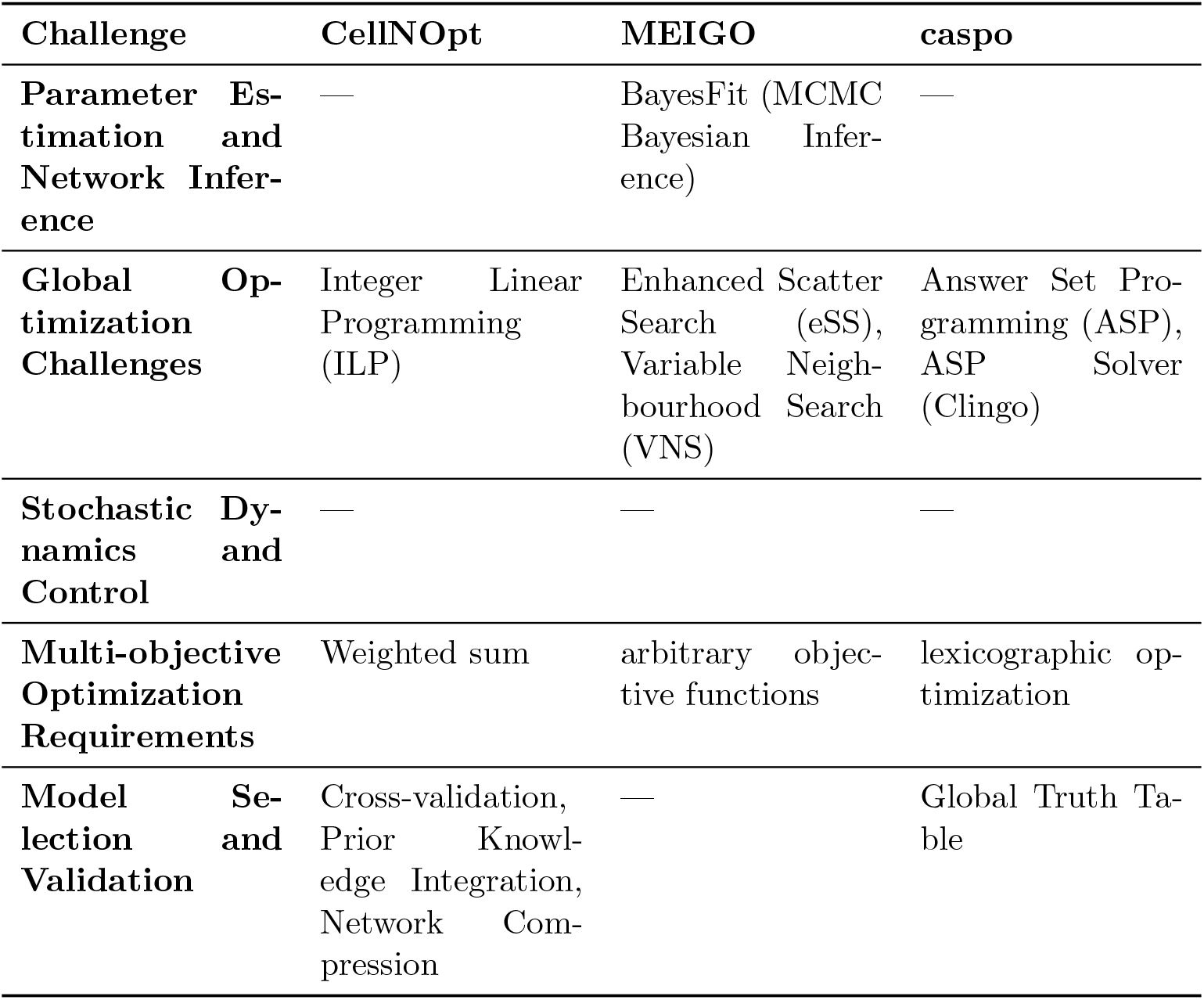
Comparative Analysis of Boolean Model Optimization Methods.

**Table 3:** Comparison of software versions available locally vs. on HPC.

| Software | Local | HPC |
| --- | --- | --- |
| Python | 3.11.11 | 3.11.5 |
| R | 4.5.0 | 4.4.1 |
| cplex studio | 2211 | 2211 |
| casq | 1.3.3 | 1.3.3 |
| rpy2 | 3.6.2 | 3.6.1 |
| caspo | 4.0.1 | 4.0.1 |
| joblib | 1.5.1 | 1.5.1 |
| networkx | 3.5 | 3.5 |
| pydot | 4.0.1 | 4.0.1 |
| pyparsing | 3.2.3 | 3.2.3 |
| scikit-learn | 1.7.1 | 1.7.0 |
| scipy | 1.16.1 | 1.16.0 |
| seaborn | 0.11.0 | 0.11.0 |
| matplotlib | 3.8.4 | 3.8.4 |
| clingo | 5.8.0 | 5.8.0 |
| SciencePlots | 2.1.1 | 2.1.1 |
| MEIGOR | 1.42.0 | 1.40.0 |
| CellNOptR | 1.54.0 | 1.52.0 |
| BoolNet | 2.1.9 | 2.1.9 |
| here | 1.0.1 | 1.0.1 |
| optparse | 1.7.5 | 1.7.5 |
| dplyr | 1.1.4 | 1.1.4 |
| tidyr | 1.3.1 | 1.3.1 |
| readr | 2.1.5 | 2.1.5 |
| data.table | 1.17.8 | 1.17.8 |

## 2 Benchmarking

Benchmarking provides essential quantitative validation of Boolean model optimization methods. This section presents a systematic framework for comparative analysis of different approaches.

The goal of this benchmark is to investigate whether structure-based optimization using experimental data leads to functionally equivalent networks in terms of dynamical behavior. A critical question we address is how many errors in prior knowledge networks can be tolerated before the derived model fails to reproduce observed dynamics.

### 2.1 Benchmark Datasets

We used three network models with associated data to benchmark the methods:

**Figure 1:**
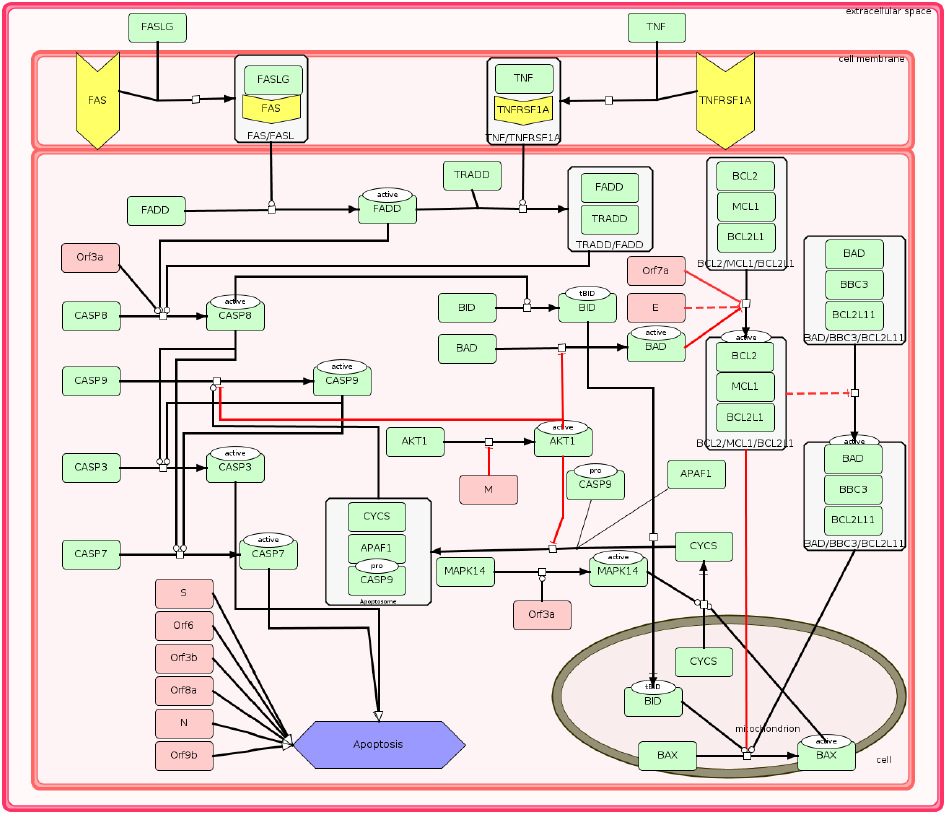
Example of a complex regulatory network: COVID-19 apoptosis submap [4].

1. **Toy dataset:** Derived from the CellNOptR R package [20], providing controlled test cases with known ground truth.
2. **DREAM challenge dataset:** Obtained from the DREAM3 challenge ^1^, offering standardized benchmarking scenarios.
3. **Other datasets:** PKNs obtained from Cell Collective ^2^ in SBML-qual format [9]. Note that ground truth experimental data are not available for these real networks.

#### 2.1.1 MIDAS Data Structure

Standardized experimental data formats are crucial for Boolean network modeling and signal transduction studies. The MIDAS (Minimum Information for Data Analysis in Systems Biology) [19] format provides a structured framework for organizing stimulus-response experimental data in tab-separated or comma-separated format.

The MIDAS structure captures complex stimulus-response relationships across multiple experimental conditions and time points through standardized column types. Each unique combination of cues (represented in TR columns) represents a specific experiment or set of interventions.

- **Treatment columns (TR:)**: Continuously active or stimulated components mirroring experimental over-expression (binary: 1 = present, 0 = don’t care)
- **Inhibitor columns (TRi:) with suffix ‘i’**: Continuously inhibited or absent components mirroring experimental knock-outs (binary: 1 = present, 0 = don’t care)
- **Data acquisition columns (DA:)**: Time step (discrete)
- **Data value columns (DV:)**: Measured protein activities or phosphorylation states with (continuous)

### 2.2 Evaluation Metrics

The comparison of optimization approaches relies on three primary categories of metrics.

- **Attractor-based metrics**: The set of attractors computed using the BoolNet [15] R package, compared using the Jaccard index, Hamming distance, and F1 score.
- **Topological metrics**: The Jaccard similarity in node and edge level.
- **Performance Metrics**: The running time and the MSE.

#### Mean Square Error

All tested algorithms internally use MSE to determine how well the current model candidate fits the experimental data. The mean squared error (MSE) is the average of the squares of the errors between the estimated values and the true value:

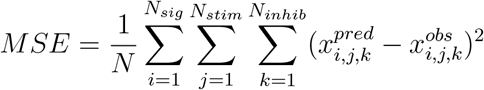

where *N* is the total number of data points, *N*_*sig*_ is the number of protein signals measured, *N*_*stim*_ is the number of stimuli, *N*_*inhib*_ is the number of inhibition conditions used between model predictions and experimental data.

The MSE computation procedure involves four key steps: simulating network states across all species and conditions, aligning simulation time points with experimental observations, excluding oscillatory or unresolved states, and calculating the mean squared error.

Oscillatory behavior detection relies on a Moving Average (MA) approach. An exponential moving average is computed and updated at each iteration using the formula:

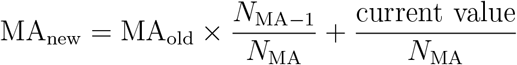

This formulation weights recent observations more heavily while preserving information from previous states. The method establishes stability thresholds where MA values outside the range [0.05, 0.95] indicate stable steady states (approaching 0 or 1), while values within this range signify oscillatory behavior. Species exhibiting oscillation are marked as *NA* (unresolved) and excluded from MSE calculations, ensuring that only converged, biologically meaningful states contribute to the model fitness evaluation.

#### Hamming distance

For two strings of equal length (or binary vectors) *a, b* ∈ {0, 1, NaN}^*n*^, the Hamming distance is the number of positions where they differ:

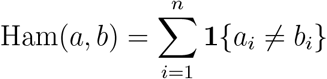

Here, it is defined only when the lengths of the strings or vectors are equal. We use Hamming distance to compare the states of all nodes in model attractors, i.e. whether a given node has the same state in attractor(s) produced by the original and candidate networks.

Note on implementation: Since attractors calculated from original and candidate networks may have different dimensions given that the algorithms may remove nodes, more sophisticated distance measures such as the longest common subsequence (LCS) or edit distance might be appropriate. However, our implementation introduces *NaN* values for missing entries, and the experimental results do not show any clear performance difference. For computational efficiency, we employ the simpler Hamming distance with *NaN* handling.

#### Jaccard similarity

For two sets *A, B* (or two binary attribute vectors), the Jaccard similarity index is

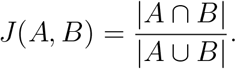

When *A, B* ∈ {0, 1}^*n*^, define the contingency counts.

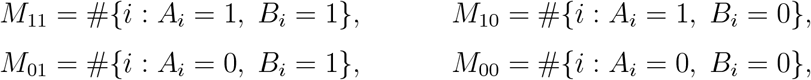

so 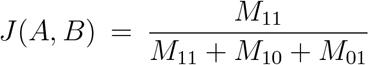 The Jaccard distance is 1 − *J* (*A, B*). In set language: *M*_00_ are elements that belong to neither set. They do not affect either the intersection or the union, so they play no role in the Jaccard similarity. If we consider *M*_00_ to also be true, Jaccard degenerates in the Hamming similarity.

#### Precision, recall and F1 score

By drawing an analogy between attractor comparison and a classification task, we can introduce analogous definitions of true positives (TP), false positives (FP), and false negatives (FN). Based on these, we can then define precision and recall in the context of attractor matching.

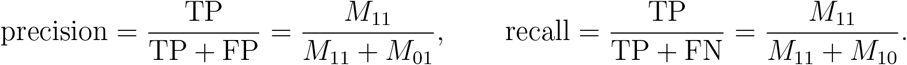

The F1 score is the harmonic mean of precision and recall:

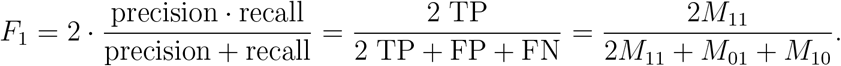

These metrics provide complementary perspectives on the performance of the optimization method, enabling a comprehensive evaluation of both topological accuracy and dynamical fidelity.

### 2.3 Benchmarking Setup

Our experimental design follows a systematic five-step validation workflow to assess the robustness of Boolean network optimization methods. This approach enables a comprehensive evaluation of how well different algorithms perform when faced with imperfect prior knowledge networks.

**Figure 2:**
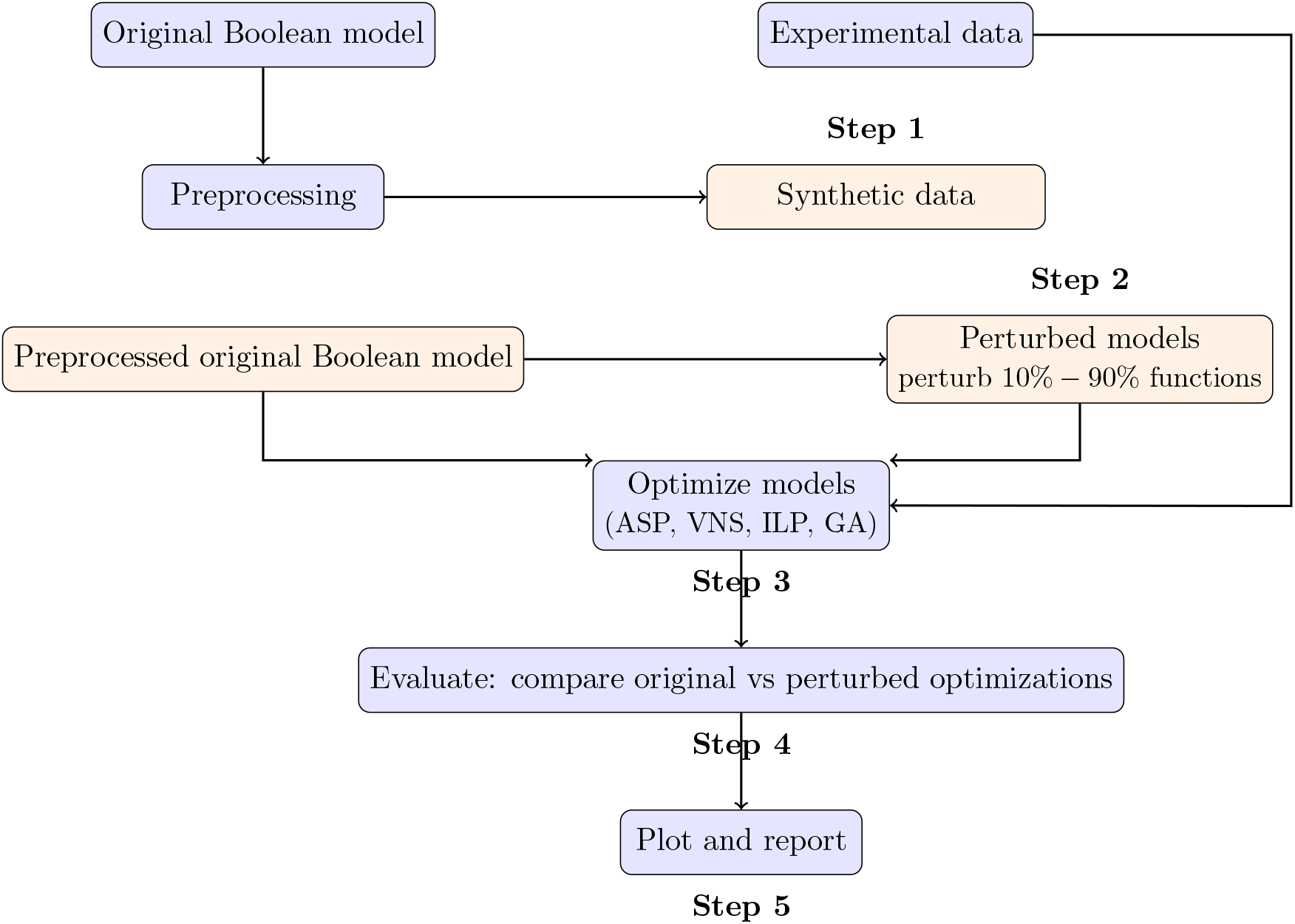
Internal validation workflow for benchmarking of tested optimization methods.

#### Step 1: Synthetic Experimental Data Generation

For data sets lacking gold standard reference datasets such as the T cell model [11] from Cell Collective, we developed a systematic approach to generate network-informed synthetic data using the BoolNet R package [15]. Our data generation pipeline employs two key functions:

1. **Experimental condition simulation:** We use *BoolNet::fixGenes* to simulate experimental perturbations by specifying which genes are over-expressed or under-expressed/knock-out, mimicking common laboratory interventions such as gene over-expression or RNA interference.
2. **Time series generation:** We apply *BoolNet::generateTimeSeries* which simulates the given Boolean network under the asynchronous updating scheme and records node states at each update cycle. We then assess how frequently each node is activated out of the entire run length, e.g. if a node has status ‘1’ 25 times out of 50 steps, we assign the value 0.5 to that node. Given the stochastic nature of the asynchronous updating scheme, we restart the simulation multiple times under the same experimental perturbation and then sample one of these runs according to the probability distribution of the returned node values. The parameters for this data synthesis are as follows:
  - numberSeries = 1000: Number of simulation restarts to generate a distribution of node states.
  - numberMeasurements = 50 or 6 *×* #Nodes:Simulation length.

#### Data processing protocol

After simulating network dynamics for a given number of time steps we sample the resulting distribution (dimensions: 1000 × #*Genes*) to extract values for the data value (DV) of each node. This approach ensures that our synthetic data capture the essential dynamical properties of the underlying Boolean network while providing sufficient statistical power for optimization method comparison.

#### Methodological Consideration

Although synthetic data cannot fully capture the complexity of real biological measurements, they maintain crucial connections to the original PKN structure. This controlled environment is sufficient for our primary objective: testing the tolerance of optimization methods to network perturbations.

#### Step 2: Modification of the Boolean network

To evaluate method robustness, we systematically introduce controlled errors into the original prior knowledge networks. We modified 10-90% of edges using four distinct modification strategies, repeating each modification level for 10 networks each, and later averaging the results. We employed four possible types of modifications:

1. **Change source/target:** Random reassignment of regulatory relationships by selecting new source and target nodes, simulating errors in interaction identification.
2. **Flip sign or direction:** For the selected edge, flip its sign (positive → negative regulation or vice versa), reverse its direction (*A* → *B* becomes *B* → *A*), or do both.
3. **Rewire:** Random reconnection of regulatory paths while preserving overall network connectivity. For example, a linear cascade *A* → *B* → *C* is transformed to *A* → *C* → *B*, maintaining the total number of connections but altering the patterns of the information flow.
4. **Split:** Division of single nodes into multiple components with appropriate edge redistribution to maintain network integrity and biological plausibility. To ensure biological realism and network functionality, we enforce several constraints:
  - **Input preservation**: Source nodes representing external stimuli remain unmodified.
  - **Connectivity maintenance**: Operations that would isolate nodes (zero incoming edges) are prohibited.
  - **Cycle prevention**: Rewiring operations that create simple two-node cycles (*A* → *B* → *A*) are forbidden to avoid trivial oscillatory behavior.

#### Step 3: Optimization

Recognizing that increased perturbation levels expand the optimization search space and computational complexity, we implemented an adaptive parameter scaling approach. The parameters of each optimization method are dynamically adjusted based on the perturbation percentage, balancing computational efficiency with solution quality.

#### Parallelization Strategy

While parallelization is essential for scaling to larger networks, several technical challenges emerge:

- **Method heterogeneity**: Runtime varies significantly among optimization approaches
- **Sequential dependencies**: Some methods, particularly integer linear programming, are challenging to implement in parallel due to reliance on third-party software
- **Cross-platform integration**: Methods implemented in different languages (R vs. Python) require careful interface management

#### Step 4: Evaluation

Perturbed networks often exhibit dramatically different dynamical complexity compared to original models. Although original networks typically have fewer than 20 attractors, optimized perturbed networks can generate more than 1000 attractors, creating significant comparison challenges.

#### Sampling-Based Comparison

For each of the 10 technical replicates per dataset, we implement a fair comparison protocol:

- **Random Attractor Selection**: Sample equal numbers of attractors from both original and perturbed networks
- **Optimal Matching**: Apply the Hungarian algorithm [12] to find optimal attractor pairings that minimize overall distance
- **Metric Computation**: Calculate evaluation metrics on matched attractor pairs

We acknowledge several limitations in our current evaluation approach. Although padding smaller attractor sets to match larger ones (creating 1000 *×* 1000 matrices) is possible, the *O*(*n*^3^) complexity of the Hungarian algorithm makes this computationally prohibitive. Investigation of attractor importance through basin size analysis revealed uniform distributions, which do not provide discriminatory power. Our results reveal inherent tensions between data-fit accuracy, graphical structure preservation, and attractor stability.

#### Computational Environment

- **Software Versions**: All methods were executed using the latest stable releases of their respective software packages, with version numbers documented for reproducibility 3.
- **Hardware Configuration**: All benchmarks were executed on AION HPC environment of the University of Luxembourg^3^.
- **Random Seed Control**: Fixed random seeds across all stochastic methods to ensure consistent comparative results.

### 2.4 Results

We conducted comprehensive experiments on three distinct datasets: Toy, DREAM, and T-Cell [11]. Each experiment was replicated 10 times to ensure statistical reliability, and the results were reported as averages across all runs.

**Figure 3:**
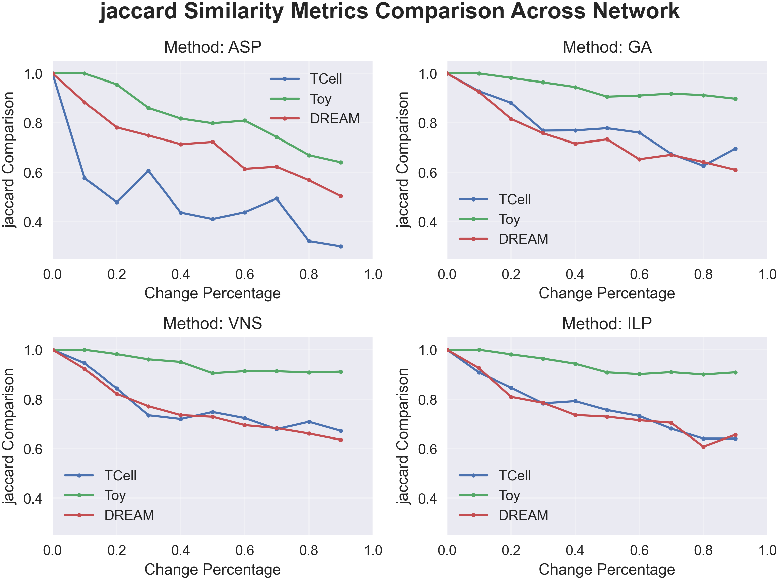
Jaccard similarity comparison between Toy, DREAM and T-Cell datasets.

**Figure 4:**
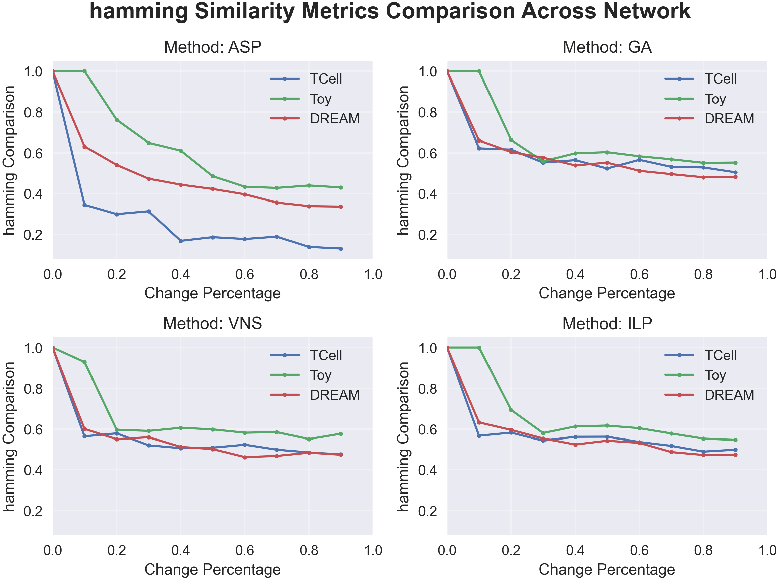
Hamming distance comparison between Toy, DREAM and T-Cell datasets.

The comparative analysis reveals a clear decreasing trend in performance as the perturbation percentage increases across all methods. Furthermore, as the size of the network increases, the performance deteriorates consistently. These trends are evident in the edge-level Jaccard similarity analysis 15, although they are not reflected in comparisons of the F1 score 16.

For Hamming similarity specifically, GA, VNS, and ILP demonstrate comparable performance characteristics. This suggests that the Hamming distance may lack sufficient sensitivity to distinguish performance differences between data sets of varying sizes.

**Figure 5:**
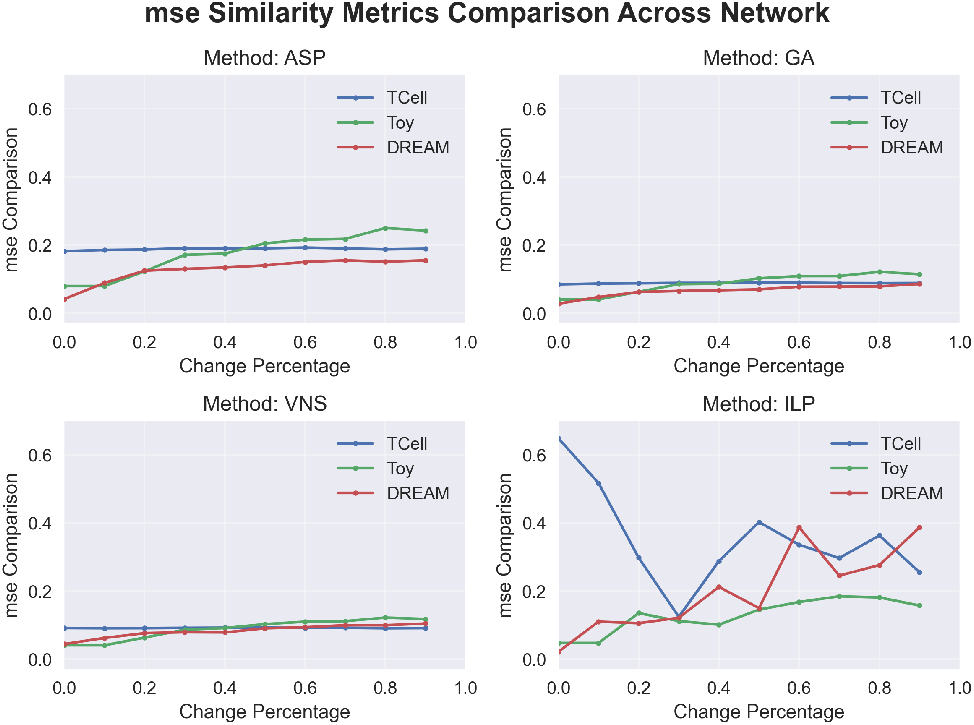
MSE comparison between Toy, DREAM and T-Cell datasets.

MSE analysis reveals several important patterns:

- ASP consistently exhibits the highest error rates across all perturbation levels
- ILP demonstrates a clear increasing trend in MSE with perturbation intensity, aligning with theoretical expectations
- ILP shows exceptionally high MSE values at zero perturbation for the T-Cell dataset, with improvement as network perturbation increases

This counterintuitive behavior in the T-Cell dataset suggests that the synthetic experimental data may not accurately represent the true structure of the prior knowledge network, implying that more testing should be done using synthetic datasets in the future.

The MSE values varied only slightly between ASP, GA, and VNS. Although the absolute scale of MSE does not fundamentally change algorithmic behavior all algorithms minimize the same loss function - it does affect sensitivity to performance differences. Small MSE variations become difficult to distinguish when all errors fall within narrow ranges (e.g., 0.15-0.25).

- **ILP (Exact Solver)**: As data becomes noisier or more perturbed, the true optimum shifts upward, and ILP accurately tracks this movement due to its exact solving capability.
- **GA**: Being a heuristic method, GA does not guarantee global optimality but explores adaptively. As the complexity of the problem increases, it occasionally finds slightly suboptimal solutions, resulting in the observed upward MSE trend.
- **VNS**: This heuristic explores neighborhoods locally and typically finds robust but approximate solutions. When data is perturbed, VNS tends to converge to similar solution types, showing less steep increases in MSE.
- **ASP**: Solutions appear to be more constrained and do not adapt as tightly to random perturbations, resulting in increasing but less steep error trends.

Mathematically, MSE can be derived as the sum of variance and bias. Perturbing the PKN (removing/adding edges, flipping logic) changes the model class over which the learning procedure searches. Our perturbation introduced both variance and bias. Removing true edges or forcing wrong edges means that the model class no longer contains the true data-generating mechanism. Therefore, bias increases (systematic error). Adding many spurious possibilities (splitting connection) can increase variance (model can overfit), and may or may not reduce bias.

Minimizing an empirical loss or variance-like term during fitting often reduces variance but does not remove structural bias introduced by a wrong PKN. In such a case, treating PKN as soft prior, regularizing the search or ensemble may mitigate the problem.

**Figure 6:**
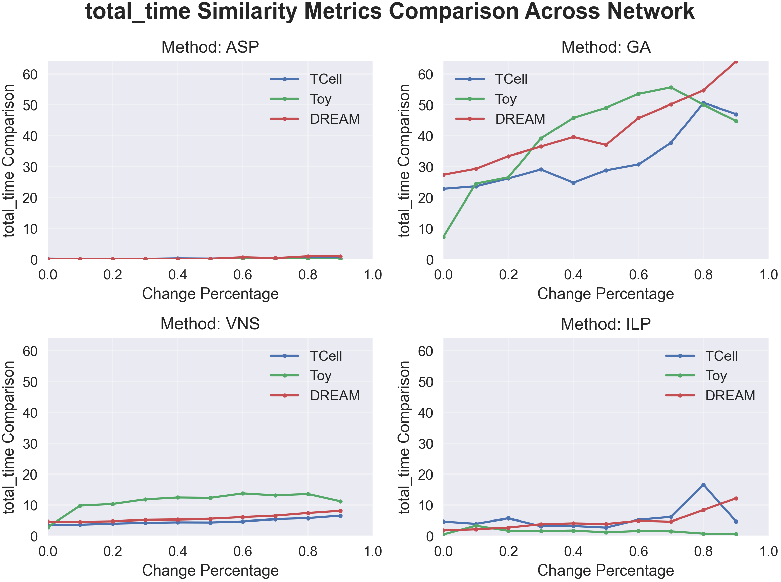
Total runtime comparison between Toy, DREAM and T-Cell datasets.

**Figure 7:**
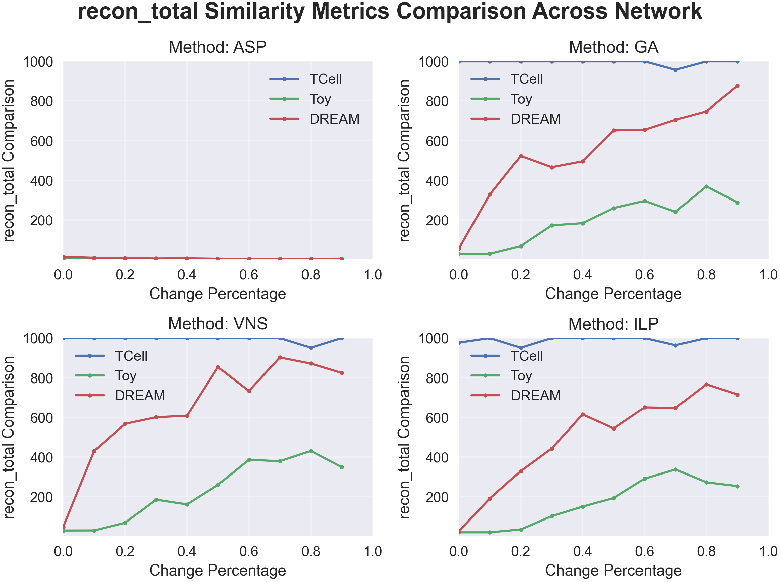
Number of reconstructed attractors: Toy vs DREAM.

Runtime analysis demonstrates significant differences in computational efficiency:

- **ASP**: Maintains the fastest execution time across all datasets
- **GA**: Requires substantially more computation time on the DREAM dataset compared to the Toy dataset, while surprisingly requiring less time for the T-Cell dataset than DREAMpotentially due to insufficient synthetic data complexity.
- **General Trend**: Increased perturbation levels generally correlate with longer optimization times.

The analysis of reconstructed attractor counts reveals that larger networks (DREAM and T-Cell) exhibit more rapid increases in attractor numbers with perturbation compared to the Toy dataset. This indicates an increase in dynamical fragmentation in larger networks, where structural perturbations more readily generate complex attractor landscapes.

#### 2.4.1 Summary of Findings

Our comprehensive benchmark analysis provides several key insights into the performance of the Boolean network optimization method.

#### Objective function

Using MSE solely as an objective function has some problems. First, minimizing MSE does not eliminate system bias. Second, MSE differences on small scales are difficult to distinguish, limiting comparative assessment. Combining complexity penalty, or multi-objective optimization to generate a Pareto frontier may mitigate this problem.

#### Scalability Characteristics

Larger and more complex networks (DREAM and T-Cell) present significantly greater reconstruction challenges both structurally and functionally. All methods exhibit performance degradation with increased network size.

#### Method-Specific Performance Profiles

ASP achieves optimal performance in topological similarity metrics while maintaining computational efficiency. ILP produces reasonable results in terms of MSE minimization, but shows a high variance. GA exhibits the longest computational requirements, but often achieves superior functional reconstruction stability. VNS shows tolerance to data perturbations.

#### Perturbation Sensitivity

All methods demonstrate critical performance thresholds around 10-30% perturbation levels, where both structural and functional reconstruction quality degrade substantially. This suggests fundamental limits to robustness in Boolean network optimization under structural uncertainty.

#### Evaluation Metric Limitations

Our analysis reveals that certain evaluation metrics (particularly the F1 score and Hamming distance) provide insufficient discriminatory power to distinguish between optimization methods, highlighting the need for more sensitive performance assessment criteria in this domain.

#### Parameter Optimization Trade-offs

The simulation step analysis reveals complex trade-offs between different optimization objectives. Extended simulations (240 steps) achieve lower MSE values, indicating better data fitting, but simultaneously reduce F1 performance and topological similarity metrics.

The experimental results collectively demonstrate that no single optimization method dominates in all evaluation criteria, suggesting that method selection should be guided by specific application requirements with respect to computational efficiency, reconstruction accuracy, and robustness to structural uncertainty. A crucial consideration of the present benchmark approach is that the systematic introduction of inaccuracies into PKNs does not reflect real-world PKN handling in BN inference. A given PKN may or may not be an accurate model of the pathway or mechanism it represents; we don’t know a priori how accurate the PKN is. As such, it is challenging to project our results on the ability of the examined methods to reconstruct BNs based on a wide variety of PKNs and datasets. Nonetheless, the present project allows us to make informed choices on the inference method and get insights into their robustness and reliability.

## 3 Future Directions

Boolean networks demonstrate strong connections to several related mathematical frameworks, particularly probabilistic graphs and neural networks. Although Boolean networks differ from causal graphs, introducing probabilistic elements (beyond stochastic processes) would enable leveraging extensive probabilistic theoretical results.

Boolean functions can be conceptualized as specialized activation functions analogous to ReLU variants:

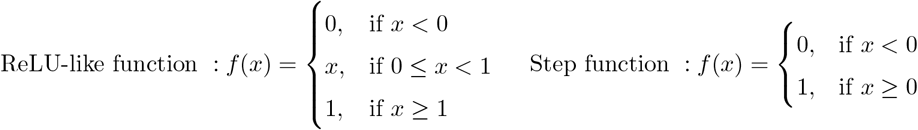

This perspective suggests that simple continuous functions could approximate complex Boolean network behaviors, potentially enabling hybrid modeling approaches. Recent work supports this view: Dersch et al. [2] introduced nary activation functions that approximate probabilistic Boolean logic using logit-based probability representations, demonstrating the ability to learn arbitrary logical ground truths within a single layer. Similarly, Cai [1] proved that a three-layer ReLU network suffices to accurately approximate any piecewise constant function.

Boolean networks may benefit from Fourier transform analysis techniques. In graph signal processing, graph signals represent mappings over network structures. The application of Fourier transforms would enable the frequency domain analysis of the dynamics of the Boolean network, similar to traditional signal processing methods. This approach could reveal periodic patterns, stability characteristics, and dynamic signatures in Boolean network behavior that are not apparent in time-domain analysis. As shown in O’Donnell’s monograph [17], any Boolean function *f*: {0, 1}^*n*^ → {0, 1} admits a unique expansion over the Boolean hypercube into parity basis functions (Fourier), where the coefficients quantify the contribution of different subsets of inputs and interaction orders. Mansour [13] further developed efficient algorithms for extracting the most significant Fourier coefficients, which is essential for handling large-scale systems. Building on this, Kesseli [10] explicitly investigated the use of Fourier spectra in the analysis of Boolean networks, demonstrating how spectral properties can provide insight into network behavior.

### Research Opportunities

Future research directions include

- Developing probabilistic frameworks for optimization Boolean Network
- Implementing graph signal processing techniques for Boolean network analysis
- Exploring approaches of using simple function approximating Boolean functions

## 4 Acknowledgments

I would like to express my sincere gratitude to Dr. Marek Ostaszewski, who provided me with this invaluable research opportunity and offered exceptional guidance throughout the entire process. I am equally grateful to Pierre Klemmer for his unwavering support, insightful feedback, and continuous encouragement. Furthermore, I extend my appreciation to Gleb Svinin for his valuable insights and engaging discussions that enriched my understanding of the subject matter. The authors acknowledge the use of ChatGPT (OpenAI) for assistance in refining language and sentence construction during manuscript preparation.

## 5 Code and Data Availability

The code and data used in this project is available on GitHub ^4^ and Zenodo.

## A Summary of Optimization algorithm

## B Software package availability

## C Result Analysis

### C.1 Individual results

**Figure 8:**
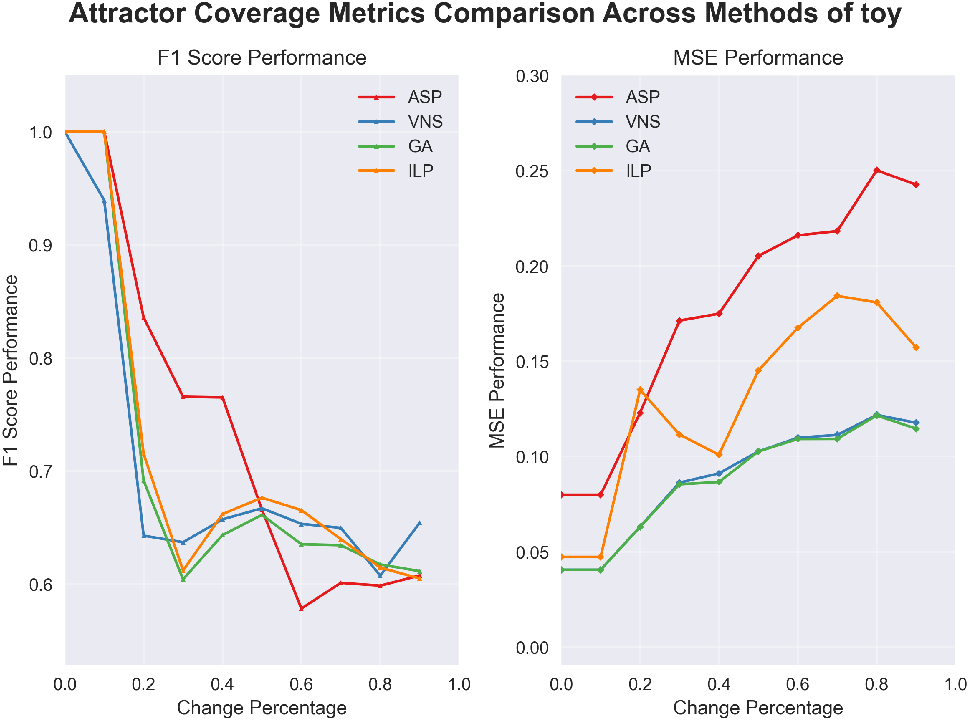
Attractor coverage metrics on the Toy dataset.

The attractor coverage analysis evaluates how frequently each optimization method (ASP, ILP, VNS, GA) successfully recovers the reference model’s attractor behavior as the prior knowledge network undergoes progressive structural perturbations. These curves serve as robustness indicators: flatter profiles indicate better preservation of dynamical outcomes under structural modifications. Our results demonstrate that ASP maintains relatively high attractor coverage at low perturbation levels but exhibits steady degradation as the perturbation intensity increases. ILP shows considerable variance across experimental replicates, while VNS and GA demonstrate comparatively stable performance in the intermediate perturbation range.

**Figure 9:**
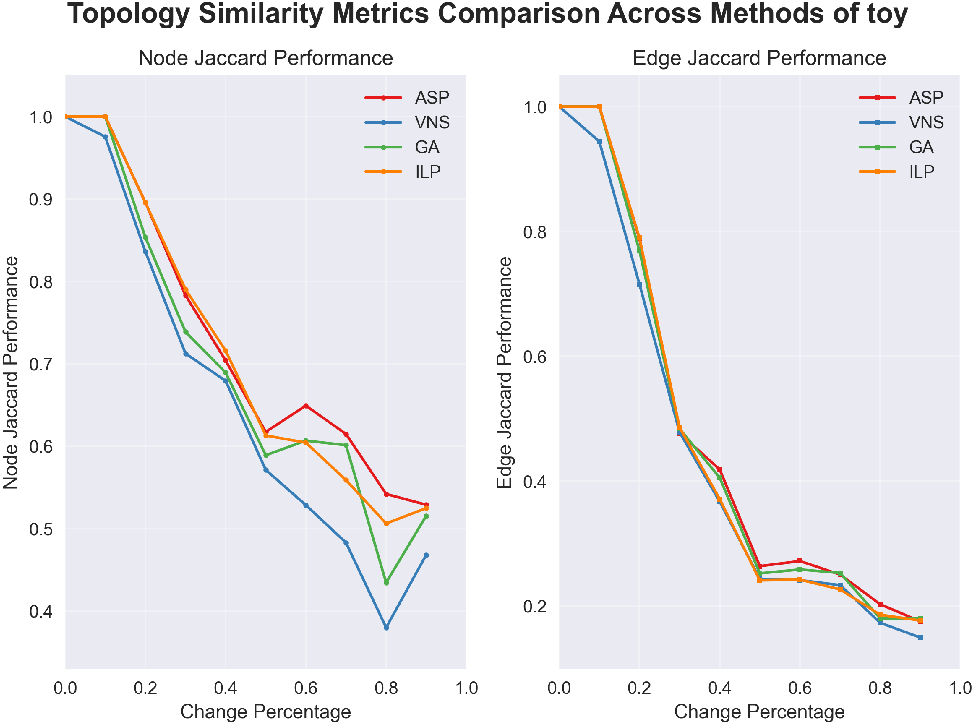
Topology-based similarity metrics on the Toy dataset.

Topological similarity metrics quantify the structural correspondence between reconstructed networks and ground truth through edge-level Jaccard indices and related graph distance measures. All methods exhibit performance degradation with increasing perturbation levels, with edge-level similarity typically declining more rapidly than coarse-grained node-level measures.

**Figure 10:**
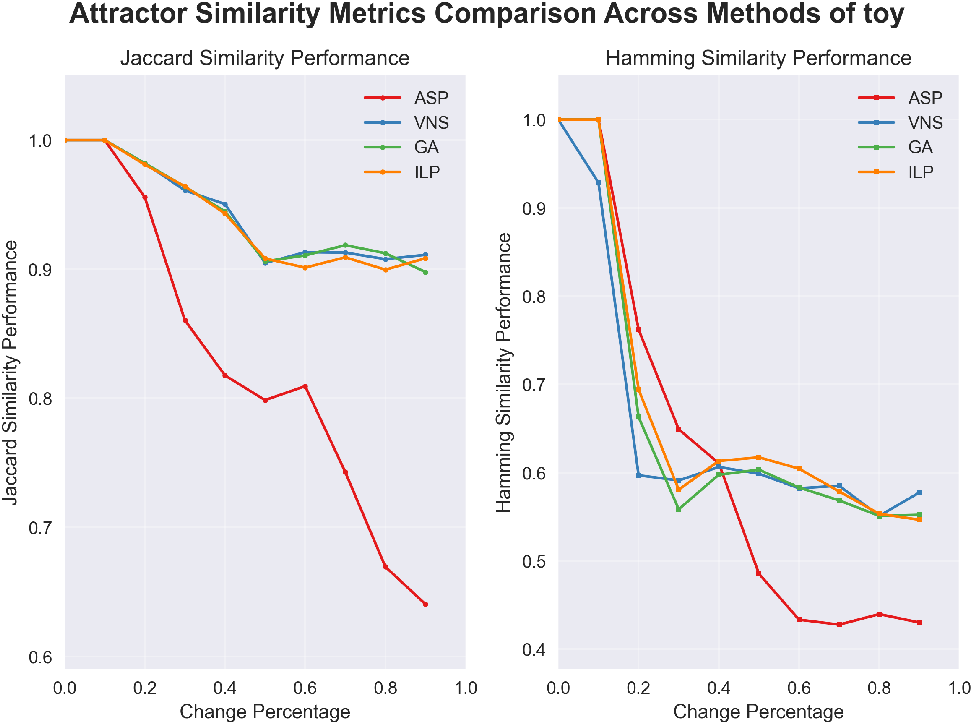
Attractor similarity metrics on the Toy dataset.

The assessment of similarity of the attractor employs multiple distance metrics (Hamming, Jaccard) to evaluate the functional correspondence of dynamical results rather than structural matching. All methods exhibit a notable performance threshold between the perturbation levels of 10 20%, indicating that moderate structural changes can substantially alter the dynamics of the system. ASP demonstrates a continuous decline with perturbation intensity, whereas heuristic methods (VNS, GA) exhibit superior functional similarity preservation at intermediate perturbation levels.

**Figure 11:**
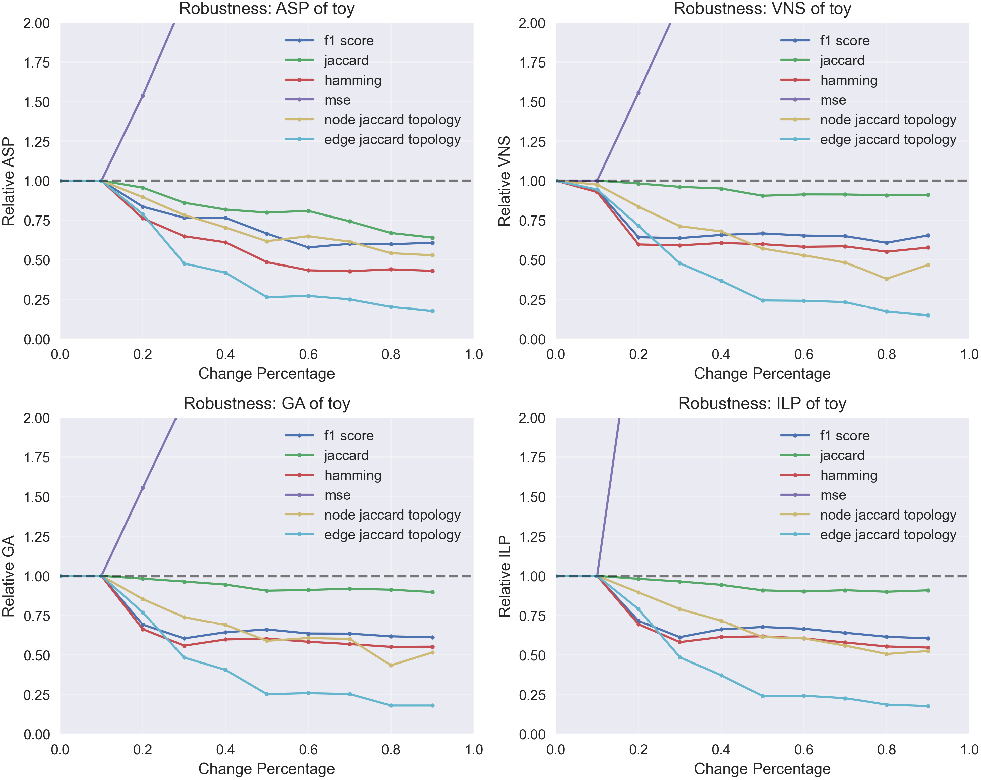
Robustness analysis on the Toy dataset.

Robustness analysis provides an integrated assessment by aggregating multiple evaluation metrics (structural and dynamical) to offer a comprehensive view of method resilience. For perturbation levels below approximately 10%, ASP, GA, and ILP maintain comparable performance characteristics. The VNS exhibits a modest performance decline in earlier perturbation stages. In particular, the edge-level Jaccard similarity decays more rapidly than others.

**Figure 12:**
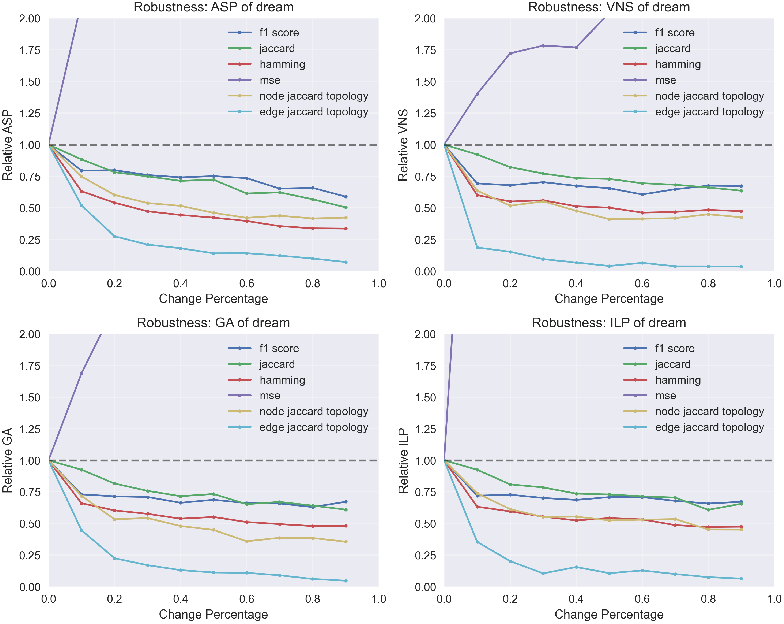
Robustness analysis on the DREAM dataset.

**Figure 13:**
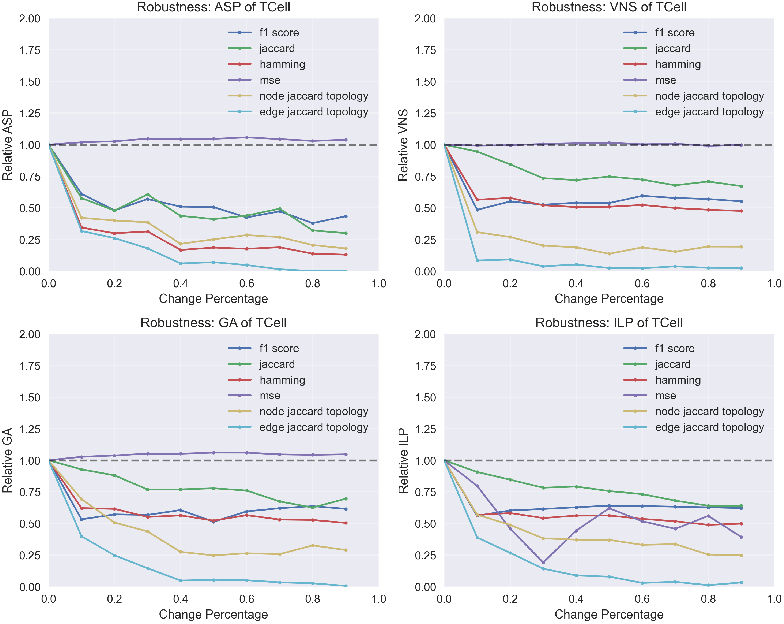
Jaccard similarity comparison between Toy, DREAM and T-Cell datasets.

The Mean Squared Error (MSE) profiles reveal distinct behavior patterns across datasets. Both the Toy and DREAM models exhibit rapid increases in MSE with perturbation level, indicating sensitivity to structural modifications. In contrast, the T-cell model displays relatively stable MSE fluctuations, which could be attributable to the synthetic nature of the experimental data used in this analysis.

**Figure 14:**
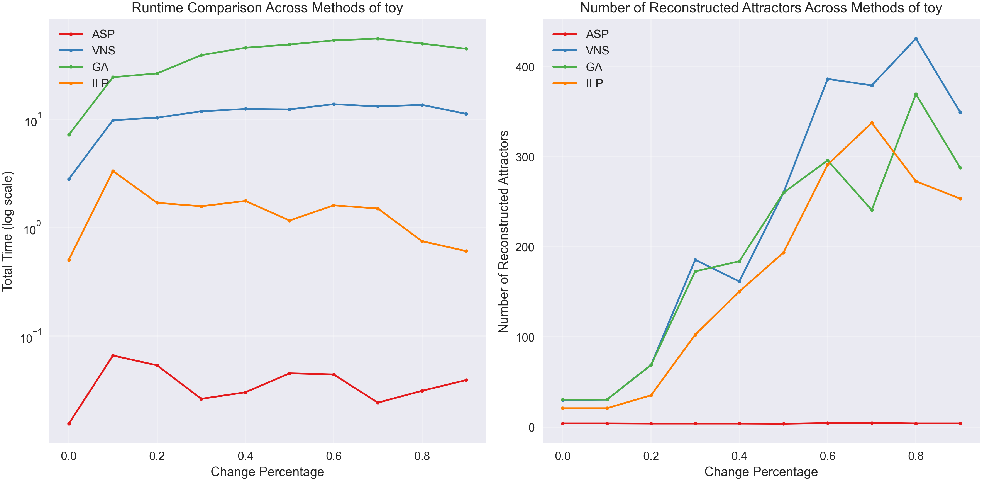
Runtime and reconstructed attractor-size comparison on the Toy dataset.

Computational efficiency and reconstruction quality analysis reveal practical trade-offs inherent in each optimization approach. ASP demonstrates the fastest execution times in these experiments while producing attractor-size distributions that closely match the reference model. GA requires the longest computational time, but often achieves more stable functional reconstructions. Other methods exhibit intermediate run-time characteristics with greater variability in reconstructed attractor counts. These results provide guidance in balancing computational cost with reconstruction quality requirements in practical applications.

### C.2 Comparison between datasets

**Figure 15:**
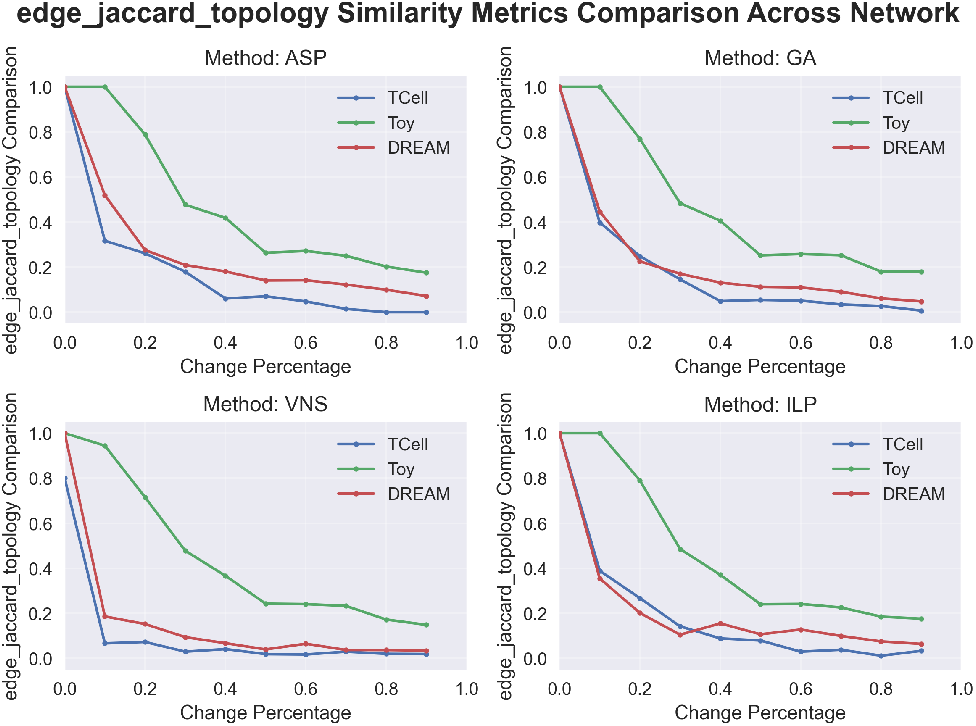
Edge Jaccard and topology similarity: Toy vs DREAM.

The relationship between network size and optimization performance is clearly demonstrated in this comparative analysis. Larger network models consistently exhibit degraded performance in all evaluation metrics. And we can say that ASP performs best in the topology sense.

**Figure 16:**
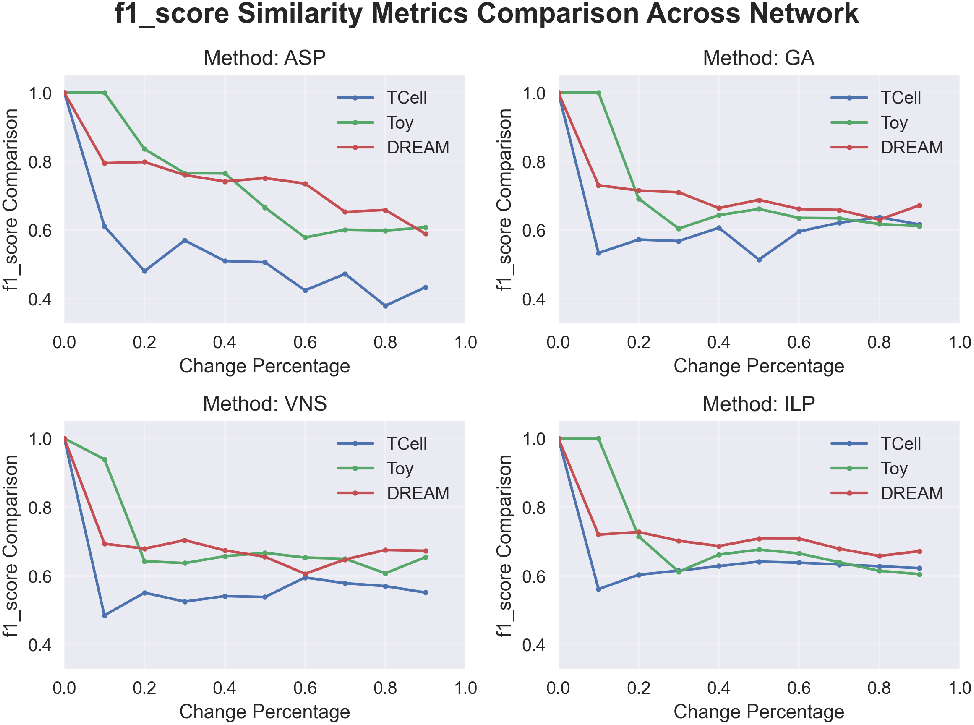
F1 score comparison between Toy, DREAM and T-Cell datasets.

The F1 score analysis fails to reveal clear relationships between network size, perturbation level, and method performance. This suggests that the F1 score may not provide sufficient discriminatory power for evaluating Boolean network optimization methods in this experimental context, as it does not effectively capture the relationship between performance characteristics and network scale or perturbation intensity.

### C.3 Parameter Analysis

**Figure 17:**
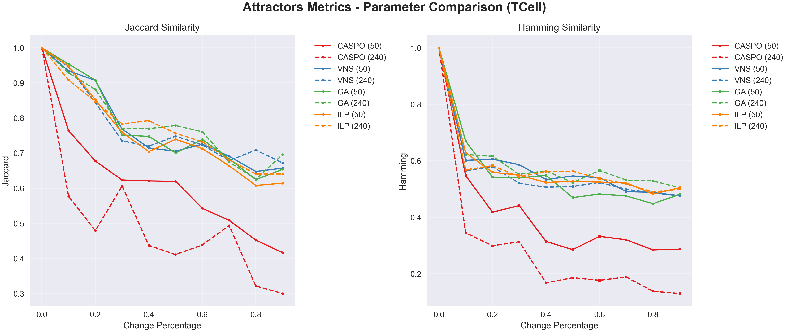
Effect of simulation steps on attractor reconstruction across optimization methods.

**Figure 18:**
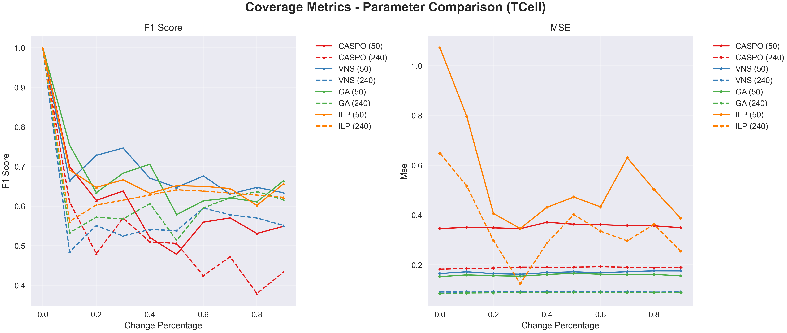
Simulation step effects on MSE and F1 performance metrics.

The analysis of simulation step parameters reveals distinct patterns across optimization methods. ASP, ILP, and GA demonstrate relatively consistent attractor reconstruction regardless of simulation length. However, ASP exhibits a counterintuitive trend in which increased simulation steps correlate with diminished performance, suggesting potential sensitivity to extended dynamics or computational overhead effects.

The right plot reveals a clear trade-off between different performance metrics. Extended 240-step simulations consistently produce lower MSE values in all methods, indicating better data fitting. In contrast, F1 performance scores decrease with longer simulations, suggesting that extended dynamics may compromise functional similarity measures. This divergence highlights the inherent tension between different optimization objectives. Moreover, 19 shows that the length of more simulations generates a more complex dynamical system.

**Figure 19:**
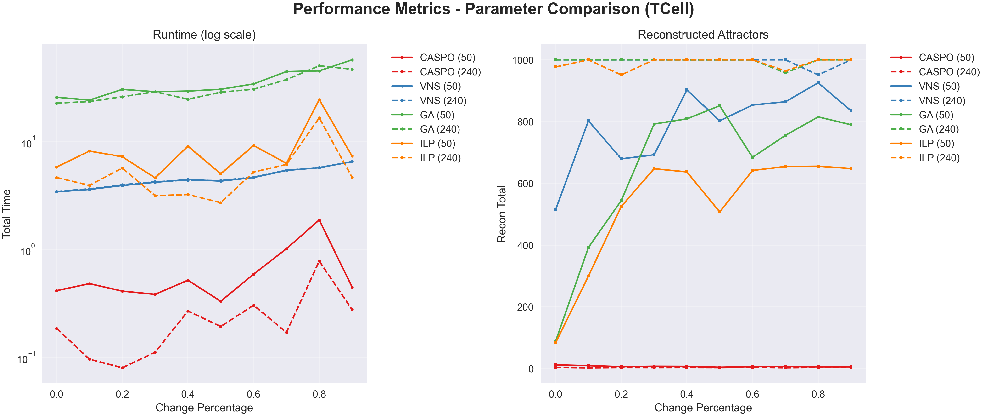
Computational complexity and dynamic system characteristics across simulation lengths.

Computational analysis reveals that 50-step simulations require more intensive optimization efforts and generate more complex dynamical systems across most methods. The sole exception is ASP, which maintains consistent computational behavior regardless of the length of the simulation. This pattern suggests that shorter simulations may capture more diverse or challenging dynamical behaviors that require greater algorithmic effort to optimize effectively.

**Figure 20:**
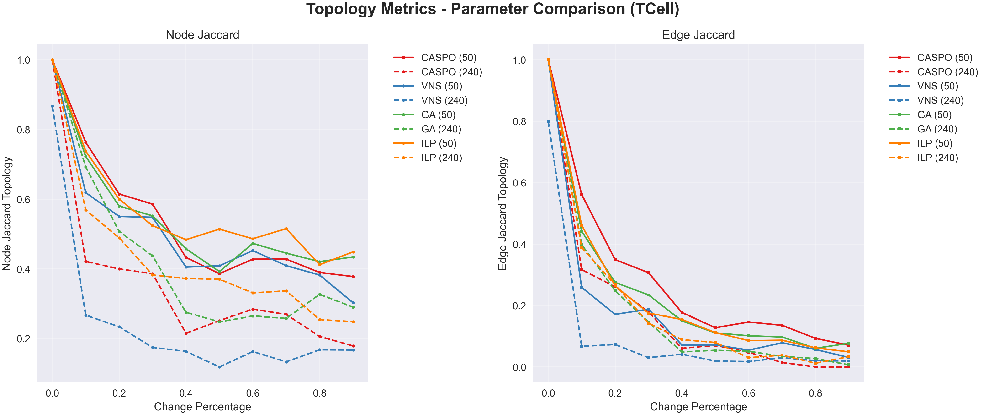
Topological similarity metrics: impact of simulation step length.

The quality of the topological reconstruction, measured through Jaccard indices at the node level and the edge level, consistently favors shorter 50-step simulations over extended 240-step protocols. This finding indicates that longer simulations may introduce dynamical complexity that obscures the underlying network structure, making topological reconstruction more challenging for optimization algorithms.

## Footnotes

1 https://gnw.sourceforge.net/dreamchallenge.html

2 https://research.cellcollective.org/

3 https://hpc-docs.uni.lu/systems/aion/

4 https://github.com/bingyulab/boolean_benchmark

## Notes

https://zenodo.org/records/17241351

https://github.com/bingyulab/boolean_benchmark/releases/tag/v1.0.0

### Competing Interest Statement

The authors have declared no competing interest.

### Summary of Updates

general revision of document format and clarifications for abbreviations and used methods across all sections Figure 2 revised.

## References

[1] Zhiqiang Cai, Junpyo Choi, and Min Liu. Relu neural network approximation to piecewise constant functions. ArXiv, abs/2410.16506, 2024.

[2] Jed A. Duersch, Tommie A. Catanach, and Niladri Das. Logical activation functions for training arbitrary probabilistic boolean operations. Information Sciences, 664:120304, 2024.

[3] Jose A. Egea, David Henriques, Thomas Cokelaer, Alejandro Fernández Villaverde, Julio R. Banga, and Julio Sáez-Rodríguez. Meigo: an open-source software suite based on metaheuristics for global optimization in systems biology and bioinformatics. BMC Bioinformatics, 15:136–136, 2013.

[4] Piotr Gawron, Marek Ostaszewski, Venkata Satagopam, Stephan Gebel, Alexander Mazein, Michal Kuzma, Simone Zorzan, Fiona McGee, Bruno Otjacques, Rudi Balling, and Reinhard Schneider. Minervaa platform for visualization and curation of molecular interaction networks. npj Systems Biology and Applications, 2:16020, 2016.

[5] Enio Gjerga, Panuwat Trairatphisan, Attila Gabor, Hermann Koch, Celine Chevalier, Franceco Ceccarelli, Aurelien Dugourd, Alexander Mitsos, and Julio Saez-Rodriguez. Converting networks to predictive logic models from perturbation signalling data with cellnopt. Bioinformatics, 36(16): 4523–4524, 2020.

[6] Carito Guziolowski, Santiago Videla, Federica Eduati, Sven Thiele, Thomas Cokelaer, Anne Siegel, and Julio Saez-Rodriguez. Exhaustively characterizing feasible logic models of a signaling network using answer set programming. Bioinformatics, 29(18): 2320–2326, 2013.

[7] Ahmed Abdelmonem Hemedan, Anna Niarakis, Reinhard Schneider, and Marek Ostaszewski. Boolean modelling as a logic-based dynamic approach in systems medicine. Computational and Structural Biotechnology Journal, 20:3161–3172, 2022.

[8] Stuart A Kauffman. Metabolic stability and epigenesis in randomly constructed genetic nets. Journal of theoretical biology, 22(3): 437–467, 1969.

[9] Sarah M. Keating, Dagmar Waltemath, Matthias Knig, Fengkai Zhang, Andreas Drger, Claudine Chaouiya, Frank T. Bergmann, Andrew Finney, Colin S. Gillespie, Tom Helikar, Stefan Hoops, Rahuman S. Malik-Sheriff, Stuart L. Moodie, Ion I. Moraru, Chris J. Myers, Aurlien Naldi, Brett G. Olivier, Sven Sahle, James C. Schaff, Lucian P. Smith, Maciej J. Swat, Denis Thieffry, Leandro Watanabe, Darren J. Wilkinson, Michael L. Blinov, Kimberly Begley, James R. Faeder, Harold F. Gmez, Thomas M. Hamm, Yuichiro Inagaki, Wolfram Liebermeister, Allyson L. Lister, Daniel Lucio, Eric Mjolsness, Carole J. Proctor, Karthik Raman, Nicolas Rodriguez, Clifford A. Shaffer, Bruce E. Shapiro, Joerg Stelling, Neil Swainston, Naoki Tanimura, John Wagner, Martin Meier-Schellersheim, Herbert M. Sauro, Bernhard Palsson, Hamid Bolouri, Hiroaki Kitano, Akira Funahashi, Henning Hermjakob, John C. Doyle, Michael Hucka, and SBML Level 3 Community members. SBML Level 3: an extensible format for the exchange and reuse of biological models. Molecular Systems Biology, 16(8):e9110, August 2020.

[10] Juha Kesseli, Pauli Rm, and Olli Yli-Harja. On spectral techniques in analysis of boolean networks. Physica D: Nonlinear Phenomena, 206(1): 49–61, 2005.

[11] Steffen Klamt, Julio Saez-Rodriguez, Jonathan A Lindquist, Luca Simeoni, and Ernst D Gilles. A methodology for the structural and functional analysis of signaling and regulatory networks. BMC Bioinformatics, 7:56, February 2006.

[12] Harold W. Kuhn. The hungarian method for the assignment problem. Naval Research Logistics (NRL), 52, 1955.

[13] Yishay Mansour. Learning Boolean Functions via the Fourier Transform, pages 391–424. Springer US, Boston, MA, 1994.

[14] Alexander Mazein, Marcio Luis Acencio, Irina Balaur, Adrien Rougny, Danielle Welter, Anna Niarakis, Diana Ramirez Ardila, Ugur Dogrusoz, Piotr Gawron, Venkata Satagopam, Wei Gu, Andreas Kremer, Reinhard Schneider, and Marek Ostaszewski. A guide for developing comprehensive systems biology maps of disease mechanisms: planning, construction and maintenance. Frontiers in Bioinformatics, 3, June 2023. Publisher: Frontiers.

[15] Christian Müssel. BoolNet–an R package for generation, reconstruction and analysis of Boolean networks. Bioinformatics, 26(10): 1378–1380, 2010.

[16] Jorge Novoa, Mnica Chagoyen, Carlos Benito, F. Javier Moreno, and Florencio Pazos. PMIDigest: Interactive Review of Large Collections of PubMed Entries to Distill Relevant Information. Genes, 14(4):942, April 2023.

[17] Ryan O’Donnell. Analysis of boolean functions. ArXiv, abs/2105.10386, 2014.

[18] Assieh Saadatpour and Rka Albert. A comparative study of qualitative and quantitative dynamic models of biological regulatory networks. EPJ Nonlinear Biomedical Physics, 4(1):1–13, December 2016. Publisher: SpringerOpen.

[19] Julio Saez-Rodriguez, Arthur Goldsipe, Jeremy Muhlich, Leonidas G. Alexopoulos, Bjorn Millard, Douglas A. Lauffenburger, and Peter K. Sorger. Flexible informatics for linking experimental data to mathematical models via datarail. Bioinformatics, 24(6):840–847, 01 2008.

[20] Camille Terfve, Thomas Cokelaer, Anake MacNamara, Dionísio Henriques, Emanuel Gonçalves, Mark K. Morris, Marc van Iersel, Douglas A. Lauffenburger, and Julio Saez-Rodriguez. Cellnoptr: a flexible toolkit to train protein signaling networks to data using multiple logic formalisms. BMC Systems Biology, 6:133, 2012.

